# Recurrent loss-of-function mutations reveal costs to OAS1 antiviral activity in primates

**DOI:** 10.1101/326454

**Authors:** Clayton M. Carey, Apurva Govande, Juliane M. Cooper, Melissa K. Hartley, Philip J. Kranzusch, Nels C. Elde

## Abstract

Immune responses counteract infections and can also cause collateral damage to hosts. We investigated functional outcomes of variation in the rapidly evolving antiviral double-stranded RNA (dsRNA) sensing factor Oligoadenylate Synthetase 1 (OAS1) in primates as a model for understanding how individual immune pathways evolve to minimize deleterious effects on host fitness. Upon binding of dsRNAs, OAS1 polymerizes ATP into 2′–5′ linked oligoadenylate (2-5A), which in turn activates Latent Ribonuclease (RNase L) to kill virus infected cells. OAS1 can undergo auto-activation by host encoded RNAs, raising the question of how it might evolve to mitigate RNase L-mediated cytotoxicity. Using a new yeast-based growth assay, we observed a pattern of frequent loss of 2-5A synthesis by OAS1 from several species. In gorillas, we identified a polymorphism in a conserved substrate binding residue that severely decreases catalytic function. In contrast, lowered 2-5A generation previously associated with variation in humans results from production of unstable OAS1 isoforms. Examination of OAS1 function in monkeys revealed a spectrum of activities, including the complete loss of 2-5A synthesis in tamarins. Frequent loss of catalytic activity in primates suggests that costs associated with OAS1 activation can be so detrimental to host fitness that its pathogen-protective effects are repeatedly forfeited.

## INTRODUCTION

The cost of immunity on organismal fitness is distributed among diverse defense functions. A growing collection of studies demonstrate that experimentally induced innate immune activation reduces longevity and fecundity^1–3^. Related work shows that increased pathogen resistance is often tied to reduced fitness in the absence of infection^4–6^. It follows that the cost-benefit balance of immune responses fluctuates between species depending on the intensity and frequency of threats from infectious microbes or changes in host biology. These differing histories of exposure can select for increased defense responses in some host lineages and decreased responses in others^7^. How these selective forces might shape the evolution of individual immune pathways remains largely undetermined. Here we investigate diversity in the antiviral Oligoadenylate Synthetase 1 (OAS1)/Latent Ribonuclease (RNase L) pathway in primates as a model system for the evolutionary balance between beneficial and detrimental outcomes of immune functions.

The collateral damage caused by OAS1/RNase L pathway activation provides a useful experimental system for studying the tradeoffs involved in evolution of immune responses. OAS proteins are a critical mediator of innate immunity and function by sensing foreign double-stranded RNA (dsRNA) from invading viruses in the cytosol^8^. Upon dsRNA binding, OAS proteins convert ATP into polymer chains joined by 2′–5′ linkage referred as oligoadenylate (2-5A)^8^. The only reported role for 2-5A is to activate RNase L, which cleaves RNAs at UA/UU dinucleotides^9^. This degradation of viral and host RNAs leads to a potent block of viral replication and eventually apoptosis in infected cells^10^. OAS1 recognizes a general motif of 17 or more base pairs of double stranded RNA with little preference for nucleotide sequence, a pattern frequently occurring in structures within the human transcriptome^11^. Indeed, constitutive editing of cellular dsRNA is required to suppress RNase L induced lethality in cultured human cells, highlighting the active measures taken to protect host cells from the deleterious effects of OAS activation in the absence of infection^12^.

Although OAS1 is the most ancient of the OAS genes, its volatile evolutionary history is consistent with potential costs. Several animal lineages, including insects and teleost fish, have lost OAS1 completely^13^. In contrast, OAS1 has undergone extensive gene amplifications in rodents and even-toed ungulates^14^. In mice, the OAS family has expanded to include eight *Oas1* genes (*Oas1a-h*). Of these OAS1 copies, only two are catalytically active, while OAS1b and OAS1d actively antagonize 2-5A generation through a dominant negative mechanism, highlighting a need to modulate harmful 2-5A levels^15,16^. OAS1 amino acid sequence is also highly variable across species. Phylogenetic and population genetic analysis demonstrate that OAS1 has evolved under recurrent positive selection in primates and other mammals, resulting in remarkable sequence diversity^17–19^. A lack of comparative studies of OAS1 activity, however, leaves the functional consequences of OAS1 evolution unknown.

Here we report that OAS1 enzymatic activity is repeatedly and independently lost or reduced among primates. In gorillas, a high frequency variant in a single active-site amino acid position leads to dramatically reduced 2-5A production. In contrast, we show that variation in human OAS1 previously associated with reduced 2-5A levels results from production of protein isoforms with lower stability rather than compromised catalytic capacity. Finally, we identify fixed amino acid substitutions shared in multiple tamarin species OAS1 that eliminate 2-5A production completely. Our findings highlight the cost-benefit conundrum of immune functions on host fitness, where deleterious effects of immune activation may sometimes outweigh the benefits of host defenses.

## RESULTS

### A yeast growth assay for OAS1/RNase L pathway activity

To investigate functional differences among OAS1 and RNase L variants, we developed an assay to efficiently measure changes in cellular viability resulting from RNase L activation by OAS1 signaling (Figure 1A). Budding yeast, which constitutively produce cytosolic dsRNA, are a proven system for studying the function of the dsRNA sensing Protein Kinase R (PKR), where eIF2α phosphorylation by PKR leads to attenuation of protein translation and growth arrest^20–22^. We tested whether heterologous co-expression of human OAS1 and RNase L might similarly arrest yeast growth due to cellular RNA degradation. While galactose-induced expression of either OAS1 or RNase L alone has no effect, coexpression of both proteins robustly inhibits yeast growth (Figure 1B). Furthermore, growth arrest specifically depends on RNase L activation, as yeast expressing OAS1 and a catalytically inactive^23^ RNase L^H672A^ grow normally (Figure 1B). Growth attenuation also corresponds with ribosomal RNA (rRNA) cleavage and reduced protein translation, a hallmark of RNase L activation^24^ (Figure 1C). Thus, OAS1/RNase L pathway function is recapitulated when genetically transplanted into yeast cells, providing a versatile new platform to efficiently measure pathway output.

**Figure 1:**
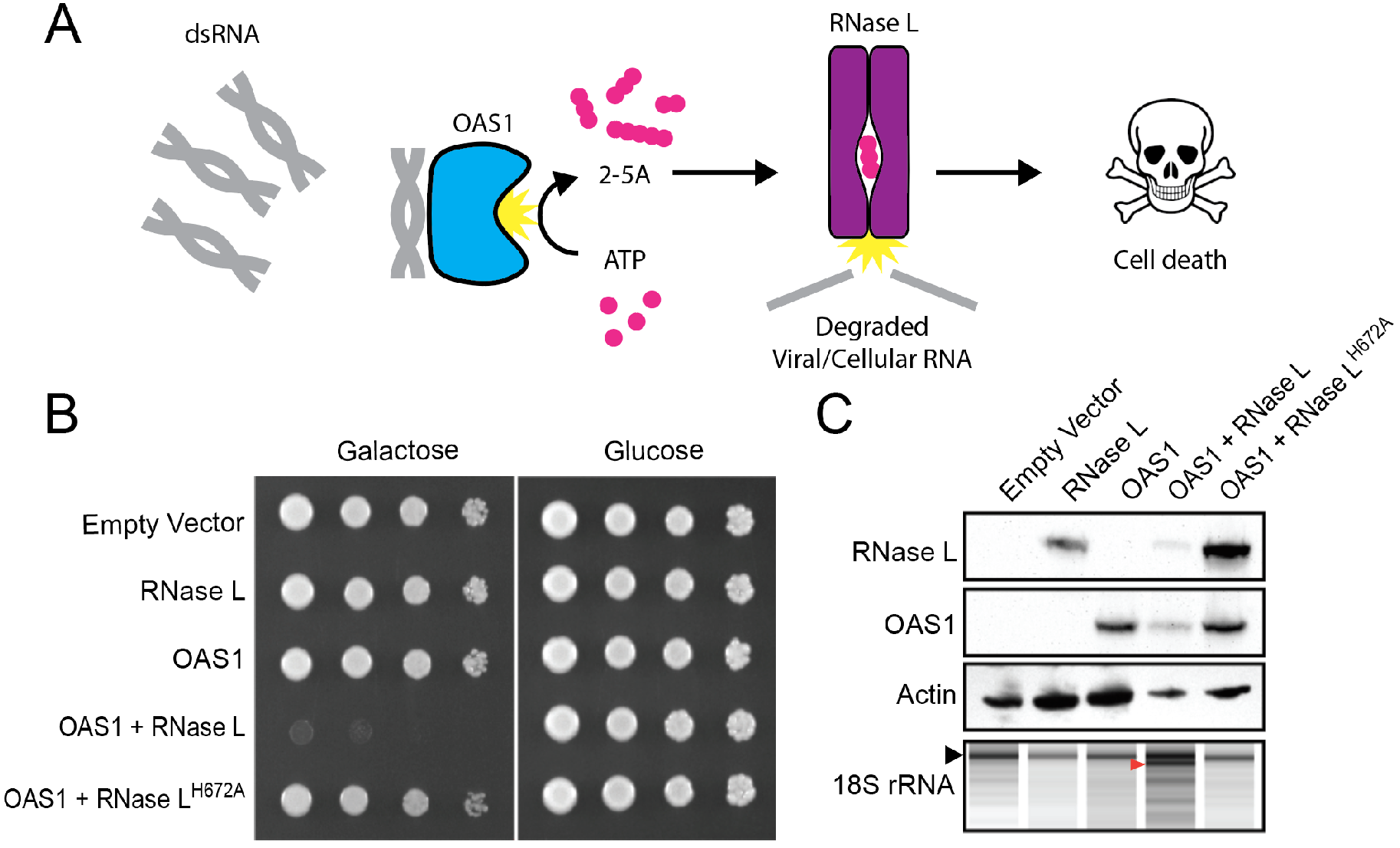
The OAS1/RNase L pathway is activated in budding yeast. (A) Schematic of the OAS1/RNaseL pathway. OAS1 is activated when bound to dsRNA, producing 2-5A chains of varying length to activate RNase L. Degradation of cellular RNAs by RNase L eventually leads to cell death. (B) Yeast strain w303 was transformed with the galactose inducible expression plasmids encoding the indicated proteins. Tenfold serial dilutions were plated on either galactose or glucose media and imaged after 24 hours (glucose) or 48 hours (galactose). (C) Immunoblot analysis of OAS1, RNase L, and actin from yeast lysates transformed with the indicated plasmids (top 3 panels). Total RNA integrity was assessed by bioanalyzer from yeast strains induced with galactose for six hours (bottom panel). Black arrow indicates 18S rRNA band, red arrow indicates RNase L cleavage products.

### A high frequency SNP in gorilla OAS1 controls catalytic output

To understand functional diversity of OAS1 in primates, we first compared the p46 isoform cloned from four hominoid species via coexpression with human RNase L in yeast. In developing the yeast assay, we found expression of the OAS1 inhibitor Us11 encoded by Herpes Simplex Virus 1 increases sensitivity and unmasks differences in OAS1 activity between species. Us11 broadly inhibits dsRNA sensing in infected cells by sequestering dsRNA^25,26^, and lowers OAS1 activity in yeast below the threshold of complete growth arrest (Figure 2A/B). Two dominant OAS1 isoforms, p42 and p46, are produced by humans and other primates and are distinguished by alternative splicing of the final exon. We initially chose to compare the longer p46 isoform to understand how variation in the C-terminus might influence activity. Unexpectedly, *OAS1* p46 mRNA cloned from chimpanzee and gorilla fibroblasts harbor early stop codons in exon five or six, respectively, resulting in smaller protein products (Figure 2A/S1B). When expressed in yeast, human, chimpanzee, and orangutan OAS1 similarly activate human RNase L and arrest growth. Yeast expressing gorilla OAS1, however, grow robustly in the presence of Us11 (Figure 2A). These data suggest that gorilla OAS1 is deficient 2-5A synthesis compared to its hominoid cousins.

To determine the basis of reduced activity in gorilla OAS1, we engineered a series of human-gorilla chimeric OAS1 proteins and tested their ability to activate RNase L in yeast (Figure S1A). With this approach, we identified an arginine to cysteine amino acid substitution at codon 130 (R130C) unique to gorilla as a candidate mutation causative of the attenuated phenotype (Figure S1B). We next tested the effects of codon 130 mutation in both human and gorilla OAS1. Reversion from cysteine 130 to the ancestral arginine in gorilla OAS1 restored activity to levels similar to other hominoids, while the converse arginine to cysteine substitution in human OAS1 significantly reduced activity as judged by robust yeast growth (Figure 2B).The R130C substitution in gorilla OAS1 is therefore responsible for its decreased level of 2-5A synthesis in yeast.

Reduced 2-5A synthesis by gorilla OAS1 might reflect reduced dsRNA binding affinity or reduced catalytic capability. To distinguish between these possibilities, we mapped residue 130 to a previously published crystal structure of OAS1 in complex with ATP analog ApCpp^27^. Consistent with a role in catalysis, Arginine 130 is located in the OAS1 active-site cleft and is highly conserved in mammals (Figure 2C). A previous structural study of porcine OAS1 identified hydrogen-bonding between residue 130 (porcine OAS1 residue 129) and the α-phosphate of the acceptor ATP molecule as a critical for proper enzymatic function^27^. To directly test the impact of the R130C mutation on catalytic function, we reconstituted OAS1 activity *in vitro* with purified proteins (Figure S3A). This technique allows for the direct visualization of 2-5A products generated from purified proteins with excess available dsRNA. Consistent with our observations in yeast, gorilla OAS1 and human OAS1^R130C^ produce far less 2-5A than wild-type human OAS1 or gorilla OAS1^C130R^ (Figure 2D). Importantly, we confirmed that gorilla RNase L functions equivalently to human RNase L in yeast and is not more sensitive to the enzymatic products of the low activity OAS1 variant (Figure S4), ruling out compensatory co-evolution. Together, these data show that the gorilla OAS1 variant recovered is catalytically impaired by a mutation in a critical active site residue.

Variation in OAS1 has previously been shown to modulate 2-5A production within human populations^28^. Extensive polymorphism also exists in nonhuman primate *OAS1*^29^, leading us to investigate whether the *OAS1* allele we recovered is fixed or variable in gorilla populations. We found evidence for polymorphism by examining a whole-genome sequencing dataset generated from 31 individual gorillas^30^. Among this cohort, the R130C allele occurred at a frequency of 36%, while the remainder encoded the full activity ancestral arginine (Table S1). The low activity allele is present in individuals from both eastern and western lowland gorilla subspecies, suggesting an ancient origin. Western gorillas are estimated to have diverged from eastern gorillas approximately 261 Kya, with some gene flow continuing between the populations until between approximately 77–150 Kya^31,32^. Therefore, genetic variation in OAS1 that dramatically reduces its catalytic output has been maintained in gorilla populations for at least 77 thousand years.

**Figure 2:**
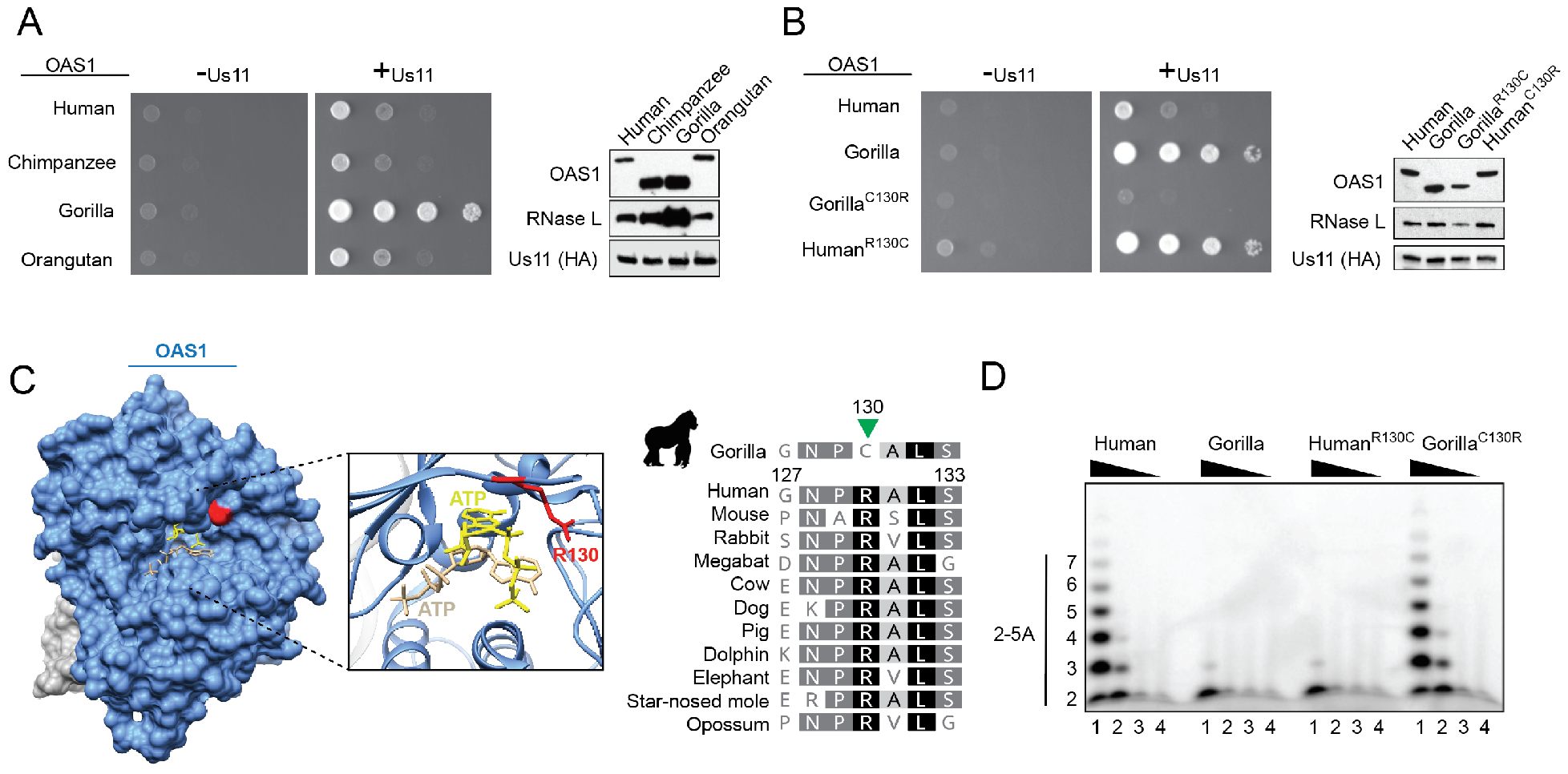
A polymorphism in gorilla OAS1 controls catalytic output. (A) Yeast constitutively expressing HSV-1 Us11 from the LEU2 locus (see methods) were transformed with vector pBM272 encoding human RNase L and OAS1 from the indicated species, plated in serial tenfold dilutions on galactose medium and imaged after 48 hours (left). Immunoblot analysis of OAS1, RNase L, and HA-tagged Us11 protein (right). (B) Procedure as in panel A, with yeast expressing gorilla OAS1^C130R^ and human OAS1^R130C^. (C) Space-filling model of crystal structure of porcine OAS1 (blue) bound to RNA (gray) in complex with ATP analog ApCpp donor (tan) and acceptor (yellow) (PDB 4RWN). Residue 130 is highlighted in red. Inset ribbon model highlighting interaction between arginine 130 and ATP (left). Amino acid sequence alignment of OAS1 residues 127–133 from a panel of mammals, highlighting substrate interacting residue 130 (green triangle) (right). (D) Gel image of ATP polymerization assay. Indicated OAS1 proteins (residues 3-346) were expressed and purified from *E. coli* and incubated with poly I:C and [a-32P] ATP for 2h before resolving by polyacrylamide gel electrophoresis (methods). Lane 1–4 are derived from reactions containing 5, 1, 0.2, or 0.04 μM OAS1, respectively.

### Variation in human OAS1 activity reflects altered isoform stability

Similar to our findings in gorillas, high frequency variation in human OAS1 have been previously associated with decreased 2-5A synthesis. A G/A SNP in the OAS1 exon 6 splice-acceptor (rs10774671) has been strongly associated with variation in human OAS activity due to alternative splicing of the final *OAS1* exon. Individuals with the ancestral G allele predominantly produce the p42 and p46 enzymes, and those with the derived A allele produce the p42, p44, p48 and p52 isoforms (Figure 3A). OAS1 activity is significantly reduced in individuals with one or two copies of the A allele^28^, leading to speculation that human isoforms differ in catalytic capacity. We investigated the mechanism of decreased 2-5A production from the p44, p48, and p52 isoforms. When expressed in yeast, both p42 and p46 robustly activate RNase L and arrest cell growth. The p44, p48, and p52 OAS1 isoforms, however, only weakly inhibit growth (Figure 3A). The expression level of these isoforms is also decreased, correlated with their reduced ability to arrest growth (Figure 3B). These results demonstrate that OAS1 isoforms associated with the A genotype are produce less stable proteins, resulting in decreased RNase L activation.

To test whether decreased expression of p44, p48, and p52 in yeast is recapitulated in human cells, we transfected HEK293T cells with expression vector encoding each isoform and measured OAS1 protein level by western blot. While p42, p46, and p48 are robustly expressed, p44 and p52 protein levels are significantly decreased (Figure 3B). Consistent with our results, a previous study failed to detect endogenous p44 and p52 isoforms in primary human cells^33^. To ensure the lack of RNase L activation is due to lowered expression, we directly tested enzymatic activity *in vitro* using equimolar amounts of full length purified protein (Figure S3B). Each of the five isoforms is capable of robust 2-5A synthesis with minimal difference between each enzyme (Figure 3C). These data indicate that reduced OAS activity in humans with the A allele is best explained by production of unstable isoforms which decrease global OAS1 protein levels and not reduced enzymatic activity of individual isoforms.

**Figure 3:**
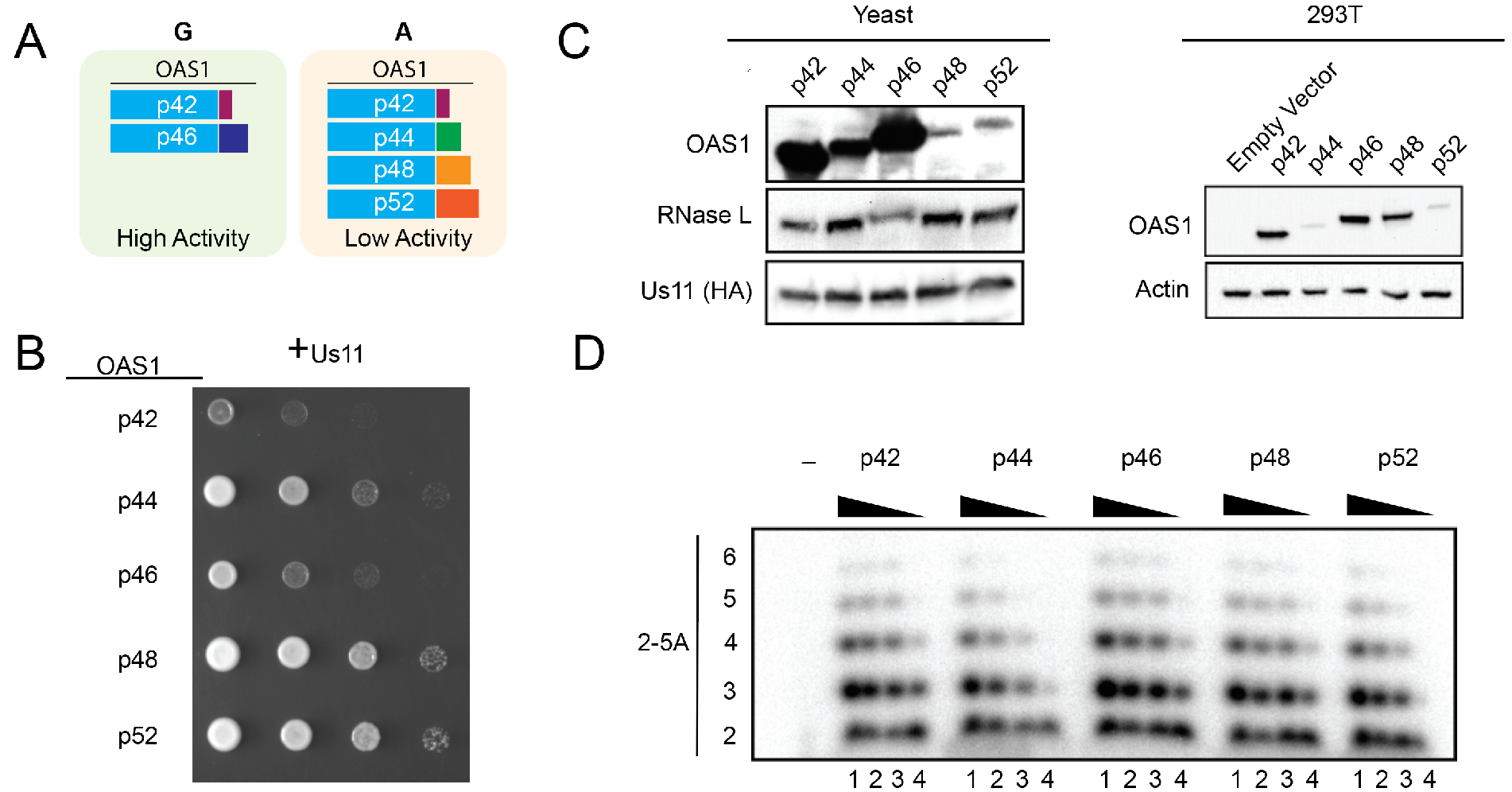
Human OAS1 isoforms differ in protein stability. (A) Summary of splice variants made by individuals with the A or G variant at exon 6 splice acceptor site (rs10774671). Individuals with the A allele have lowered OAS activity (see text). (B) Tenfold serial dilutions of yeast expressing human RNase L and the indicated human OAS1 isoform imaged after 48 hours of growth on galactose medium. (C) Immunoblot analysis of OAS1, RNase L and Us11 expression from yeast (left) or OAS1 and Actin from HEK293T cells transfected with 3ug expression vector encoding the indicated OAS1 isoform (right). (D) Gel image of ATP polymerization assay. Indicated human OAS1 isoforms were expressed and purified from *E. coli* and incubated with poly I:C and [α–32P] ATP for 2h before resolving by polyacrylamide gel electrophoresis (methods). Lane 1–4 are derived from reactions containing 5, 1, 0.2, or 0.04 μM OAS1, respectively.

### OAS1 activity is variable among monkeys and lost in tamarins

To determine if loss-of-function variation in human and gorilla reflects a general pattern in OAS1 evolution, we next tested OAS1 function from a more diverse sampling of primate species. To capture a broad taxonomic and genetic distribution, we cloned *OAS1* p46 from five Old World and three New World monkeys and tested their ability to activate human RNase L in yeast. Similar to gorilla, dusky titi *OAS1* p46 contains an early stop codon in exon 6, resulting in a smaller protein product (Figure S1). African green monkey, rhesus macaque, baboon, and dusky titi OAS1 robustly activate human RNase L and inhibit yeast growth. Both talapoin and white faced saki OAS1, however, display an intermediate phenotype with lowered relative levels of RNase L activation (Figure 4A). Consistent with reduced yeast growth inhibition, *in vitro* 2-5A synthesis assays confirmed that talapoin OAS1 produces less 2-5A compared to baboon or African green monkey OAS1 (Figure 4B). In contrast, yeast expressing saddleback tamarin OAS1 grew robustly, even in the absence of Us11, indicating a complete loss of OAS1 enzymatic function (Figure 4A/5B). OAS1 activity is therefore highly variable among the monkey species sampled in our study, with alleles ranging from high, intermediate, or zero capacity to activate RNase L in yeast.

**Figure 4:**
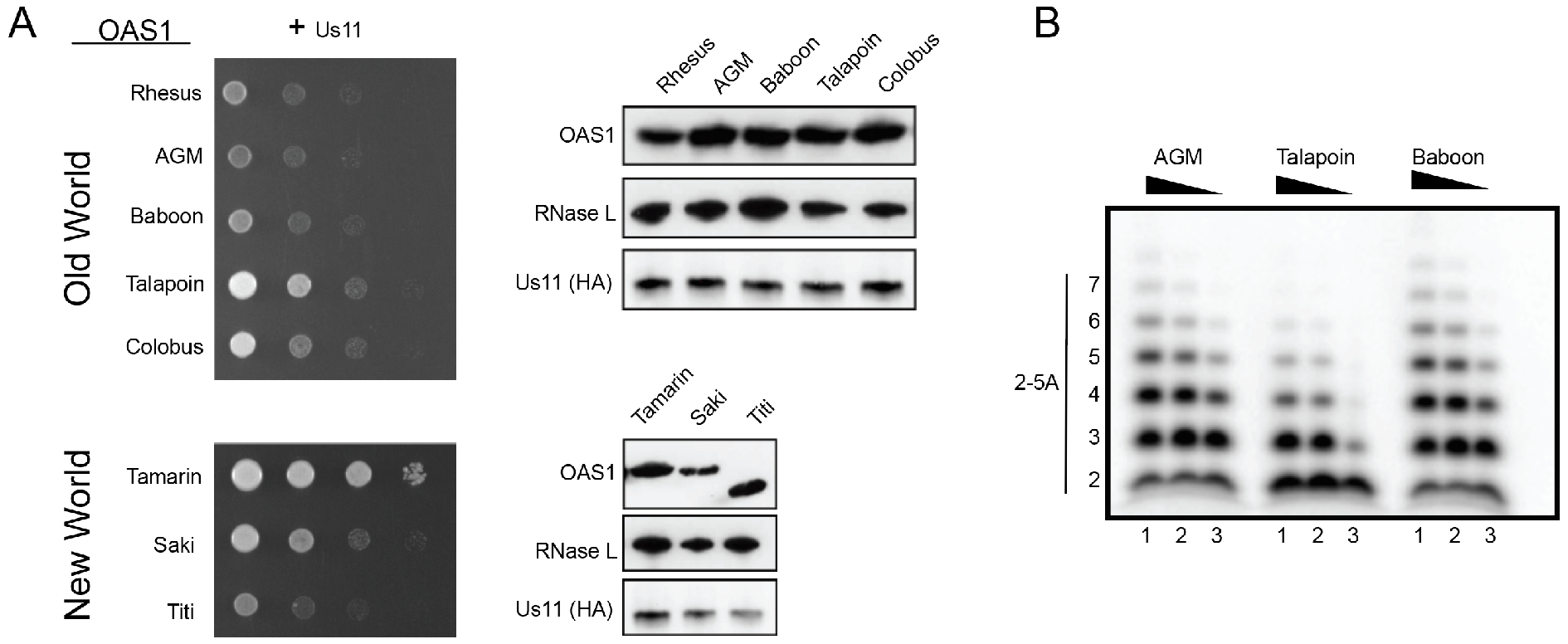
Variable levels of OAS1 activity among New World and Old World monkeys. (A) Tenfold serial dilutions of yeast expressing human RNase L and OAS1 from the indicated species (AGM – African green monkey) were imaged after 48 hours of growth on galactose medium (Left). Immunoblot analysis of OAS1, RNase L and HA-Us11 from yeast cell lysates after 6h of galactose induction (Right). Dusky titi OAS1 p46 is truncated by a stop codon in exon 6 and produces a smaller protein product. (B) Gel image of ATP polymerization assay. Indicated OAS1 proteins (residues 3–346) were expressed and purified from *E. coli* and incubated with poly I:C and [α–32P] ATP for 2h before resolving by polyacrylamide gel electrophoresis (methods). Lane 1–3 are derived from reactions containing 5,1, or 0.2 μM OAS1, respectively.

To determine the basis of the loss of 2-5A synthesis by saddleback tamarin OAS1, we first performed pairwise amino acid alignments to identify possible mutations in conserved active-site residues. A short helical turn motif (p-loop) plays an important role in catalysis through an interaction with the substrate nucleotides^11,27^. Remarkably, saddleback tamarin OAS1 has three mutations in p-loop residues otherwise strictly conserved in mammals (S63F, S74I and D77N) (Figure 5A). Notably, residue 77 is one of three catalytic aspartates that coordinate with Mg^2+^ions to facilitate catalysis. In addition, serine 74, which hydrogen bonds with the β-phosphate of the donor ATP, is replaced with a hydrophobic phenylalanine in tamarin. Finally, like the hypomorphic gorilla OAS1 allele, tamarin OAS1 independently acquired amutation at residue 130 (arginine to histidine), additionally altering its interaction with the acceptor ATP. Thus, saddleback tamarin OAS1 contains multiple debilitating mutations in conserved substrate interacting and catalytic residues critical for 2-5A synthesis.

To chart the evolutionary history of OAS1 loss-of-function in tamarins, we PCR amplified and sequenced OAS1 exons 1–6 from genomic DNA of the closely related mustached tamarin. Mustached tamarin shares mutations in noncatalytic residues (S63F, S74I, and R130H) but has an intact aspartate triad, similar to a partial sequence available from the related Midas tamarin^34^ (Figure 5A). We next tested whether mustached tamarin OAS1 retains residual 2-5A synthesis activity compared to saddleback tamarin. Yeast expressing saddleback or mustached tamarin OAS1 and human RNase L, however, display no growth arrest(Figure 5B), indicating a lack of activity by both species. Furthermore, purification (Figure S3C) and visualization of enzymatic activity *in vitro* revealed no 2-5A production above background from either species (Figure 5C). Finally, we investigated whether tamarin OAS1 might preferentially activate tamarin RNase L through another mechanism. Although saddleback tamarin RNase L also displays lowered activity than human RNase L, it is not activated by tamarin OAS1 in yeast (Figure S5). Thus, evolution of OAS1 in the tamarin lineage involved several inactivating mutations in conserved substrate coordinating residues 63, 74, and 130 in a common ancestor, and the subsequent loss of catalytic aspartate 77 in saddleback tamarins.

**Figure 5:**
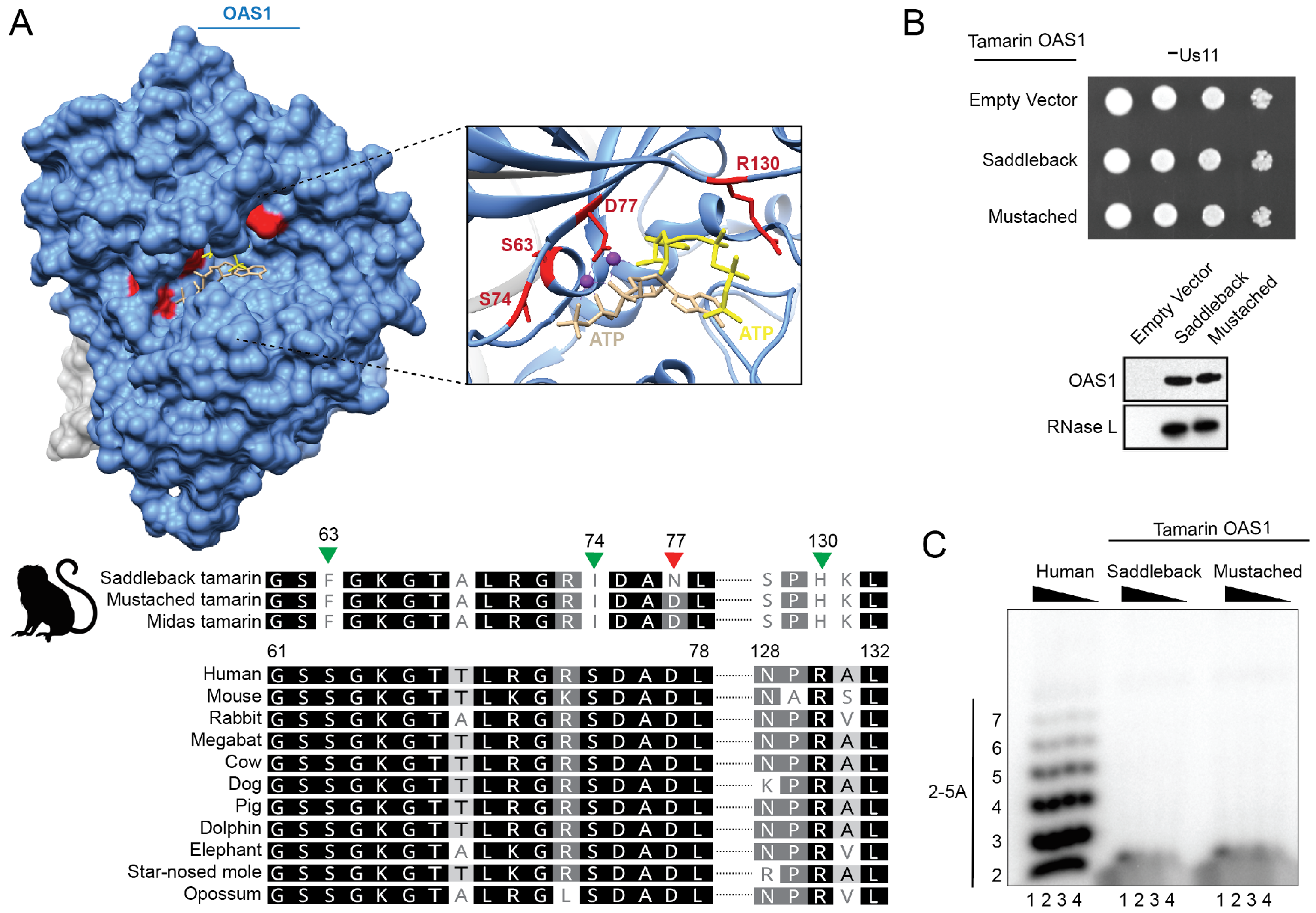
Complete loss of catalytic activity in tamarin OAS1. (A) Space-filling model of crystal structure of porcine OAS1 (blue) bound to RNA (gray) in complex with ATP analog ApCpp donor (tan) and acceptor (yellow), and Mg^2+^ (purple) (PDB 4RWN). Conserved residues 64, 74, 77, and 130 mutated in tamarin OAS1 are highlighted in red (top). Amino acid sequence alignment of OAS1 residues 61–78 and 128–132 from the indicated mammalian species. Green triangles indicate mutations in conserved residues implicated in substrate interaction, red triangles indicate mutations in catalytic residues (bottom). (B) Serial dilutions of yeast strain w303 transformed with empty integrating pRS405 vector at the LEU2 locus and pBM272 encoding human RNase L and mustached or saddleback tamarin OAS1 were grown on galactose medium and imaged after 48 hours of growth (left). Immunoblot analysis of tamarin OAS1 and human RNase L expressed in yeast (right). (C) Gel image of ATP polymerization assay. Indicated OAS1 proteins (residues 3–346) were expressed and purified from *E. coli* and incubated with poly I:C and [α–32P] ATP for 2h before resolving by polyacrylamide gel electrophoresis (methods). Lane 1–4 are derived from reactions containing 5, 1, 0.2, or 0.04 μM OAS1, respectively.

## DISCUSSION

Robust immune defenses provide clear benefits to host fitness yet can also exact great costs. Our discovery of repeated and independent debilitating mutations in primate OAS1 reveals distinct mechanisms for balancing the antiviral benefits and deleterious costs of 2-5A synthesis. These findings also highlight how evolution of innate immunity can be constrained by deleterious outcomes of cytotoxic defense mechanisms.

Previous studies on the impact of altering the OAS/RNase L pathway in healthy and infected animals revealed contrasting outcomes consistent with evolutionary compromises. Mice lacking RNase L exhibit greatly increased susceptibility to infection with multiple viruses^35,36^, supporting a fundamental role for the OAS/RNase L pathway in immunity. In the absence of infection, however, RNase L deficient mice display significantly increased longevity, potentially due to the lack of chronic low level activation of OAS proteins by cellular dsRNAs^37^. Indeed, RNA extracted from various cell types weakly activates OAS1 *in vitro^38^,* consistent with detectible 2-5A in uninfected tissues^39^. In addition, recent work demonstrated that cellular RNAs activate OAS proteins in the absence of RNA editing by ADAR1^12^. Lineage-specific increases in the repertoire of cellular RNAs that activate OAS1 could therefore account for the benefit of forfeiting 2-5A synthesis in some species. OAS2 and OAS3, which have distinct dsRNA recognition patterns^40^, provide additional means of RNase L activation, potentially alleviating the loss of OAS1 activity during infection.

Despite recovering several *OAS1* loss-of-function alleles in this study, we did not find species lacking the potential to encode OAS1, for example through mutations resulting in premature stop codons precluding the production of the core enzyme. Recurrent, and even compound, mutations in critical substrate binding and catalytic residues in otherwise intact proteins is consistent with a role for OAS1 beyond 2-5A synthesis. Our findings support an evolutionarily conserved role for RNase L-independent antiviral functions for OAS1 that have been proposed in other studies^41,42^.

Genes encoding immune functions are rich with genetic variation in humans^43^. In some cases, common genetic variants in key mediators of cellular immunity lead to partial or complete loss of protein function and can also be associated with disease. The human OAS1 variant associated with lower *in vivo* 2-5A levels is also associated with increased risk of infection with West Nile virus^44^ and development of Sjörgens syndrome^33^, an autoimmune disease associated EBV and CMV infection. Despite these risks, the allele resulting in production of unstable OAS1 isoforms rose to near fixation in ancient African populations, proceeded by reintroduction of the ancestral high activity allele via introgression from Neandertals to non-African populations^45–47^. Our identification of multiple independently evolved loss-of-function OAS1 alleles in nonhuman primates is consistent with a common underlying evolutionary pressure frequently favoring lowered OAS activity in humans and nonhuman primates alike.

Similar pressures might favor loss of function in other immunity genes as well. For example, a common human variant of STING, a central component of several viral DNA sensing pathways, results in decreased IFN-β production in response to viral DNAs^48,49^ (but also see^50^). The recent description of dampened STING signaling in bats further highlights how selection can favor variants that decrease immune responses to accommodate host biology in specific lineages^51^. Taken in isolation, the prevalence of alleles compromising defense functions is difficult to explain. When considered in a broader context, our findings illustrate how evolutionary forces can act on individual proteins to repeatedly select for diminished immune functions across species.

## METHODS

### Cloning of OAS1 and RNase L cDNAs

Total RNA was isolated from primate cell lines (quick-RNA miniprep, Zymo Research) followed by preparation of cDNA (Superscript III First-Strand Synthesis kit, ThermoFIsher scientific). Unless otherwise noted, cell lines were obtained from Coriell Cell Repositories (Figure S1; Genbank accession: MH188449-MH188461). OAS1 and RNase L cDNAs were PCR amplified from cDNA using primers indicated in Table S2. Mustached tamarin OAS1 p46 exons 1–6 were PCR amplified and sequenced directly from genomic DNA. The assembled mustached tamarin OAS1 coding sequence was synthesized by Life Technologies. The human OAS1 p44 and p52 splice variant sequences were synthesized by Life Technologies based on sequences deposited in genbank. OAS1 and RNase L variants were cloned into the yeast CEN Gal10/Gal1 dual expression plasmid pBM272. Human OAS1 splice variants were cloned into vector pcDNA6 for mammalian cell expression. OAS1 was cloned via EcoRI/NotI restriction sites, and RNase L through BamHI sites introduced by PCR primers. C-terminally hemagglutinin tagged Us11 (Us11-HA) was codon optimized for budding yeast and synthesized by Life Technologies (based on Genbank YP_009137147.1.

### Yeast growth assays

Yeast strain w303 was used in all experiments in this study. Standard techniques were used for culturing and transformation of yeast. Strains were generated by transformation of EcoRV linearized vector pRS405 containing either an empty site (-Us11) or HA-Us11 (+Us11), resulting in integration at the leu2 locus. Yeast strains were transformed with pBM272 containing OAS1/RNase L variants. Transformants were grown in S -leu -ura liquid media containing 2% glucose overnight and washed before plating. Tenfold serial dilutions were prepared (OD_600_ 3.0, 0.3, 0.03, 0.003) and a multichannel pipette was used to plate 2.3 μl of each dilution on 2% glucose or 2% galactose medium. Unless otherwise noted, glucose plates were imaged after 24 h of growth, galactose plates after 48 h of growth.

### RNA integrity analysis

Transformed yeast were grown to log phase in 2% raffinose S -leu -ura growth medium, followed by induction with 2% galactose for 6 h. Total RNA was extracted using a YeaStar RNA kit (Zymo research) and quantified. RNA integrity was assessed using an RNA ScreenTape assay with the Agilent TapeStation instrument.

### Western Blotting

Transformed yeast were grown to log phase in 2% raffinose S -leu -ura growth medium, followed by induction with 2% galactose for 6 h. Cell lysates containing equal amounts of yeast were prepared by treatment with 0.1 M NaOH for 5 min, followed by lysis in 2⨯ SDS loading buffer. Mammalian cell lysates were prepared by direct lysis in 2× SDS buffer containing 8 M Urea and 3 M thiourea. Total protein was resolved by Mini-PROTEAN GTX polyacrylamide gel electrophoresis (Bio-rad). Proteins were detected using anti-OAS1 (Sigma HPA003657, 1:1000), anti-RNase L (Santa Cruz sc-23955, 1:200), anti-HA (Covance MMS-101P, 1:1000), and anti-Actin (BD 612657, 1:1000) antibodies. Blots were visualized using film or a Li-C or c-digit chemiluminescent imager.

### Mammalian OAS1 alignment

The following OAS1 coding sequences were retrieved from genbank for multialignment: human (*Homo sapiens*, NM_016816.3), mouse (*Mus musculus*, NM_145211.2), rabbit (*Oryctolagus cuniculus*, XM_017349609.1), megabat (*Pteropus Alecto*, NM_001290162.1), cow (*Bos Taurus*, NM_001040606.1), dog (*Canis lupus familiaris*, NM_001048131.1), pig (*Sus scrofa*, NM_214303.2), dolphin (*Lipotes vexillifer*, XM_007468577.1), elephant (*Loxodonta africana*, XM_010598964.2), starnosed mole (*Condylura cristata*, XM_012730965.1), opossum (*Monodelphis domestica*, XM_001378771.3). Translated sequences were aligned with cloned tamarin or gorilla OAS1 sequences using clustalw.

### Human cell culture and transfection

5 × 10^5^ 293T cells in 6-well plates were transfected with 3 μg of expression vector pcDNA6 (Invitrogen) encoding each OAS1 isoform or an empty site using FuGENE HD (Promega) transfection reagent according to the manufacturers specifications. Cells were harvested 48 h post transfection for western blot analysis.

### Generation of Chimeric and mutated OAS1 proteins

Human-gorilla OAS1 chimeras were generated using PCR stitching. Overlapping PCR products were generated and gel purified. 50ng of each PCR product was then added to a new PCR reaction and allowed to anneal and extend for 12 cycles before addition of primers to generate full length chimeric *OAS1* coding sequences. Plasmids encoding human OAS1^R130C^’ gorilla OAS1^C130R^, and RNase L^H672A^ were generated by site directed mutagenesis.

### Primate OAS Isoform Purification

Human and other primate OAS isoforms were PCR amplified and cloned into a custom pET vector optimized for expression of an N-terminally tagged 6×His-MBP fusion protein, and proteins were expressed and purified from BL21(DE3) RIL *E. coli* as previously described ^52^. Briefly, competent cells already containing the pRARE2 tRNA plasmid (Agilent) were transformed with OAS expression plasmid and grown in 2 × YT media at 37°C to an OD_600_ of ~0.6 before cooling on ice for 15 min, induced with 0.5 mM IPTG and incubated with shaking at 16°C for ~18 h. Cell pellets were re-suspended in lysis buffer (20 mM HEPES-KOH pH 7.5, 400 mM NaCl, 10% glycerol, 30 mM Imidazole, 1 mM DTT), lysed by sonication, and protein was isolated from clarified lysate using Ni-NTA resin (Qiagen) and gravity-flow chromatography. Resin was washed with lysis buffer supplemented to 1 M NaCl to remove bound nucleic acid, and MBP-OAS protein was eluted with lysis buffer supplemented to 300 mM Imidazole. MBP-OAS proteins were diluted to 50 mM Imidazole and 5% glycerol and concentrated to ~30 mg ml^−1^ prior to overnight digestion with Tobacco Etch Virus protease at 4°C. Digested protein was separated from MBP and protease using a 5 ml Ni-NTA column (Qiagen) and subsequently purified by Superdex 75 16/60 size-exclusion chromatography in gel-filtration buffer (20 mM HEPES-KOH pH 7.5, 250 mM KCl, 10% glycerol, 1 mM TCEP-KOH). Purified OAS was concentrated to ~10 mg ml^−1^, flash-frozen in liquid nitrogen and stored at −80°C for biochemical experiments. MBP-tagged human OAS1 splice isoforms and MBP-tagged mustached tamarin and saddleback tamarin OAS homologs were purified using identical conditions, except proteins were dialyzed into storage buffer (20 mM HEPES-KOH pH 7.5, 250 mM KCl, 10% glycerol, 1 mM TCEP-KOH) overnight at 4°C immediately following Ni-NTA elution and directly stored without tag removal or gel-filtration purification.

### OAS Enzymatic Assay

Purified recombinant OAS enzyme was incubated in reaction buffer (10 mM Tris-HCl pH 7.5, 25 mM NaCl, 10 mM MgCl_2_, 1 mM DTT, 0.1 mg ml^−1^ BSA) supplemented with 200 μM ATP and [α-^32^P] ATP (~2 μCi). 10 μl reactions included final OAS enzyme concentrations as indicated, or enzyme titration with final concentrations of 0.04, 0.2, 1, and 5 μM, and included ~5 μM poly I:C (Invivogen) for dsRNA stimulation. Reactions were incubated at 37°C for 2 h and terminated by adding an equal volume of stop solution (95% deionized formamide, 20 mM EDTA) and incubation at 95°C for 2 min. Reactions were analyzed by denaturing gel electrophoresis on a 32-cm tall 20% polyacrylamide 7 M urea gel with 0.5 × TBE running buffer, and 2′–5′ oligoadenylate products were visualized using a Typhoon phosphoimager (GE Healthcare).

## ACKNOWLEDGEMENTS

The authors thank Jennifer Doudna (UC-Berkeley) and Harmit Malik (Fred Hutchinson Cancer Research Center) for reagents and support during initial experimental setup. We also thank to Mark Hochstrasser (Yale University) for providing us with the double-gal, dual expression plasmid pBM272. We thank Tom Sasani for assistance with analysis of gorilla population genetic data. We thank Dustin Hancks and Florian Maderspacher for valuable feedback on the manuscript. N.C.E. is supported by the NIH (R01GM114514, P50GM082545), the Burroughs Wellcome Fund Investigators in the Pathogenesis of Infectious Disease Program and is a H.A. & Edna Benning Presidential Endowed Chair (University of Utah). C.M.C. is supported by the Microbial Pathogenesis Training grant (NIH T32AI055434). P.J.K. is funded by the Claudia Adams Barr Program for Innovative Cancer Research and Richard and Susan Smith Family Foundation. A.A.G. is supported by a US National Science Foundation Graduate Research Fellowship.

---

**Figure S1:**
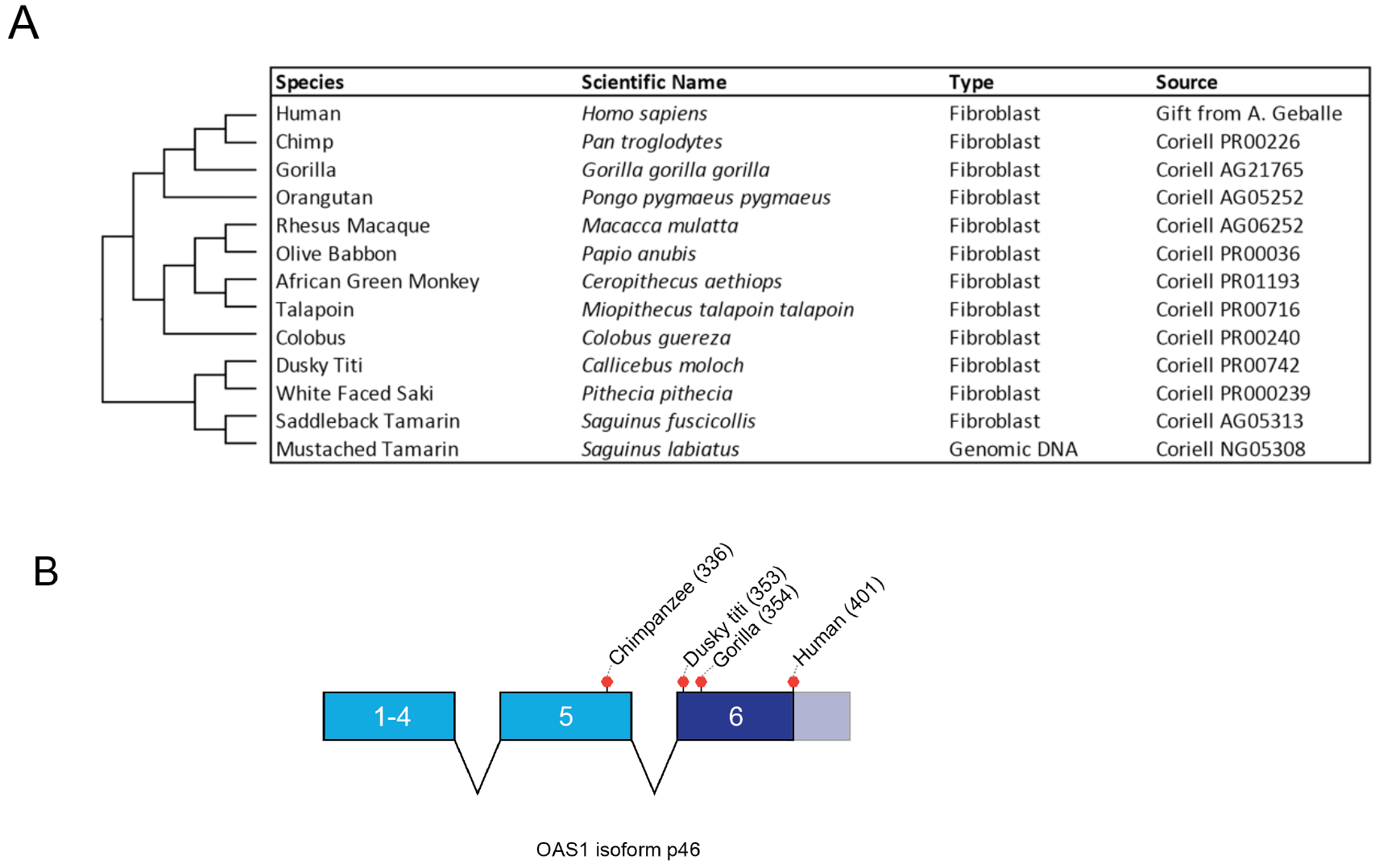
Source of primate OAS1/RNase L. (A) Source material used for cloning OAS1/RNase L cDNA and genomic DNA. (B) Schematic of OAS1 exons 1–5 and alternatively spliced exon 6 found in isoform p46. Premature stop codon positions are indicated for chimpanzee, dusky titi, and gorilla *OAS1* p46 cloned from cDNA.

**Figure S2:**
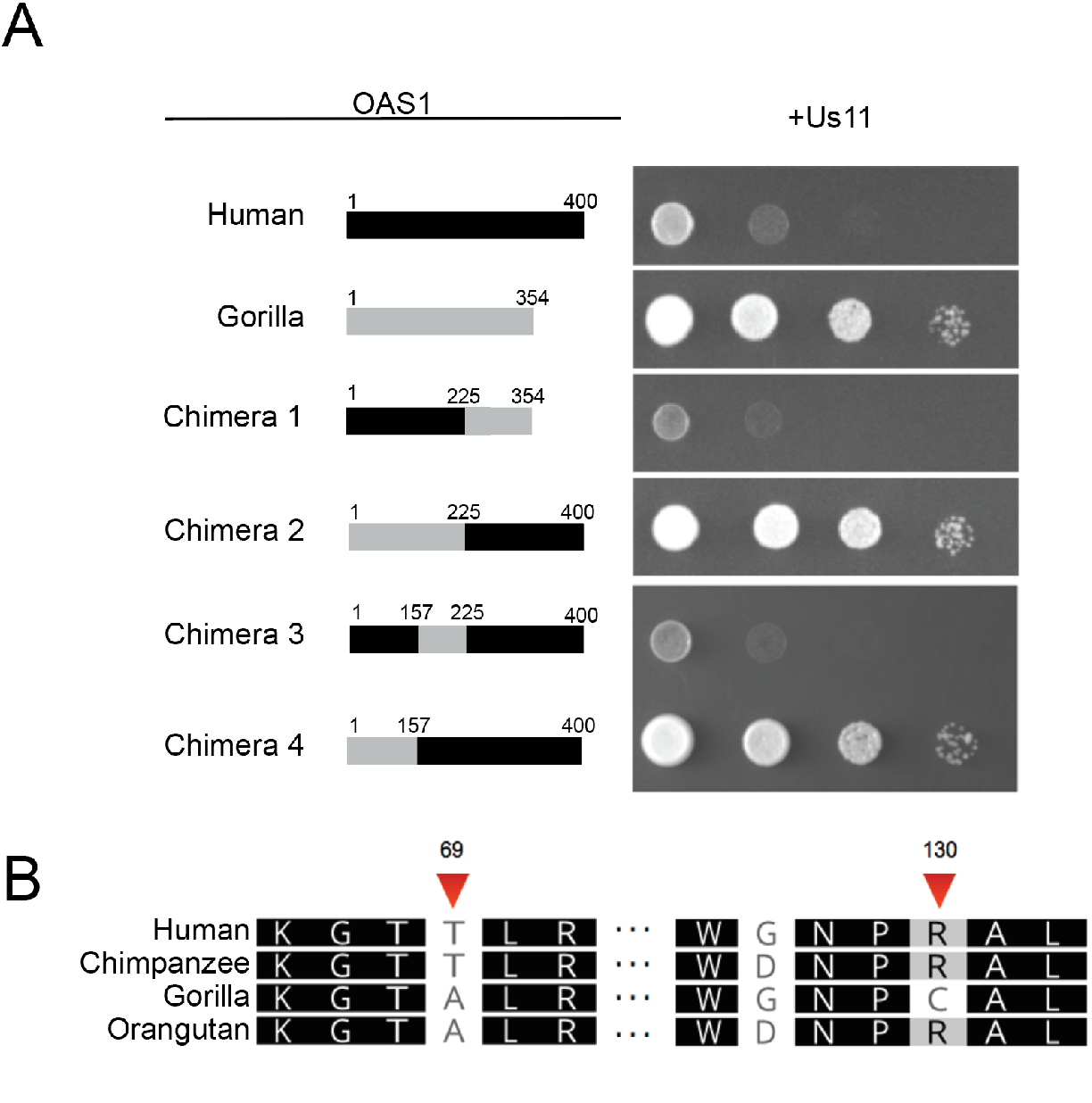
A substitution in gorilla OAS1 between amino acid 1–157 confers lowered activity. (A) Yeast strain w303 constitutively expressing HSV1 Us11 transformed with human RNase L and the indicated OAS1 proteins. Chimeras were constructed by overlapping PCR amplification (methods) using either human (black) or gorilla (gray)OAS1 templates. Amino acid positions at chimera boundaries are indicated. (B) Amino acid alignment of hominoid OAS1 highlighting the two amino acid differences between human and gorilla OAS1 between positions 1–157.

**Figure S3:**
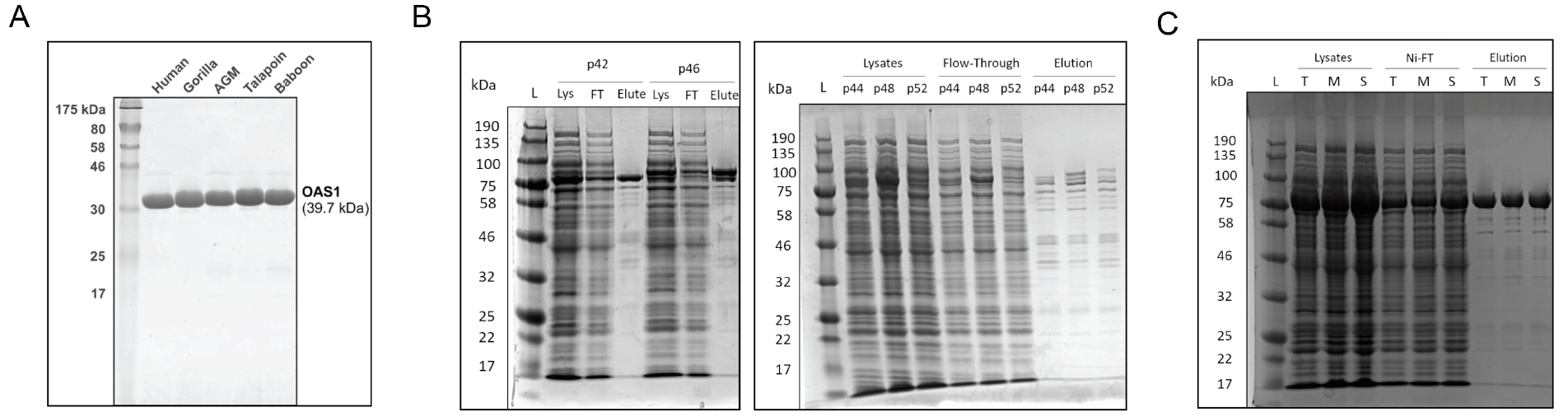
Purification of OAS1 proteins. (A) Indicated OAS1 proteins (3–346; ending near the truncating stop observed in several primates) after purification and protease cleavage of his-MBP tag. (B) Purification of full length human OAS1 isoforms before protease treatment (Lys = *E. coli* cell lyate, FT = Ni column flow through, Elute = Ni column elution). (C) Purification of tamarin OAS1 proteins (3–346) (S = Saddleback tamarin, M = mustached tamarin, T = identical to S).

**Figure S4:**
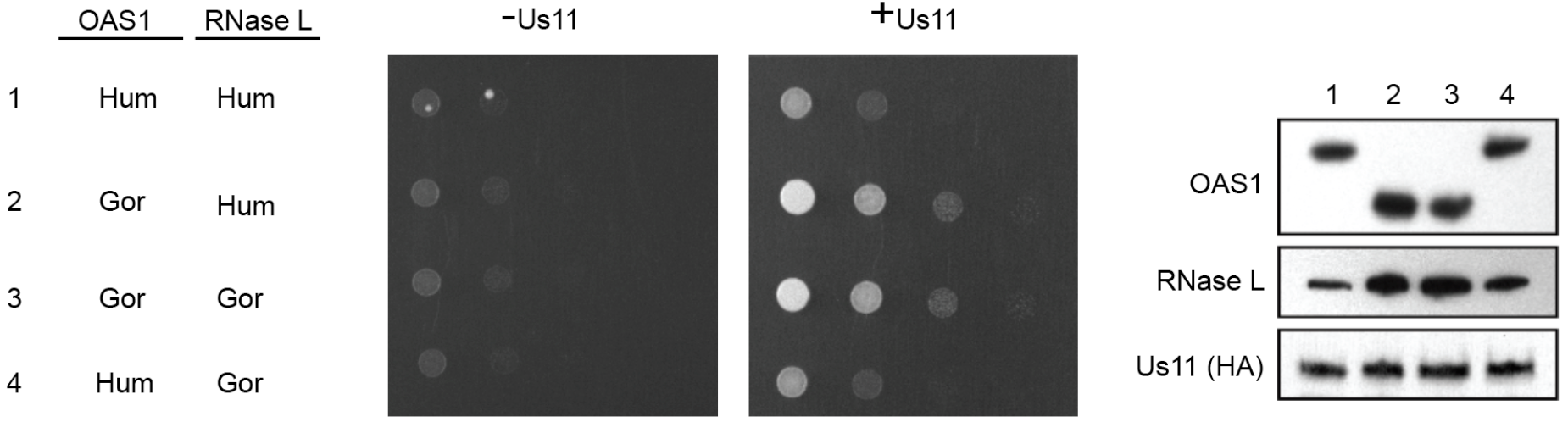
Gorilla RNase L shows no preference for gorilla OAS1. Yeast strain w303 expressing HSV-1 Us11, OAS1 and RNase L from the indicated species imaged after 24 hours of growth on galactose medium (Hum - Human, Gor - Gorilla) (right). Immunoblot analysis of OAS1, RNase L and HA-Us11 from yeast induced with galactose for 6 hours (left).

**Figure S5:**
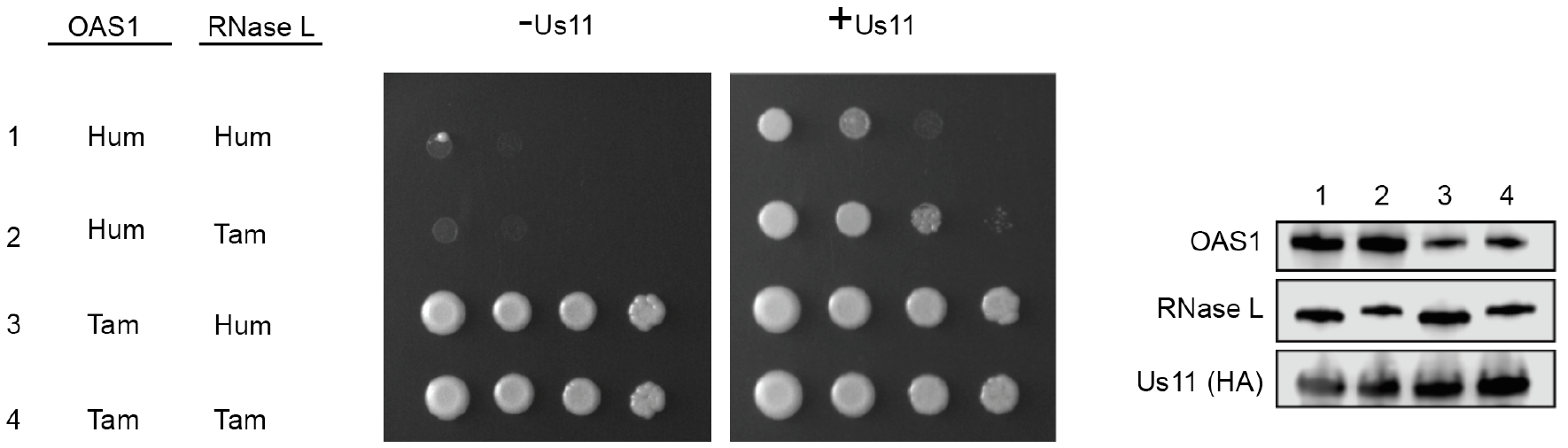
Tamarin OAS1 does not activate human or tamarin RNase L. Yeast strain w303 expressing HSV-1 Us11, OAS1 and RNase L from the indicated species (Hum - Human, Tam - saddleback tamarin) imaged after 72 hours of growth (right). Immunoblot analysis of OAS1, RNase L and HA-Us11 from yeast induced with galactose for 6 hours (left). Decreased signal from tamarin OAS1 may be due to lower expression level or lowered antibody reactivity.

**Table S1.**
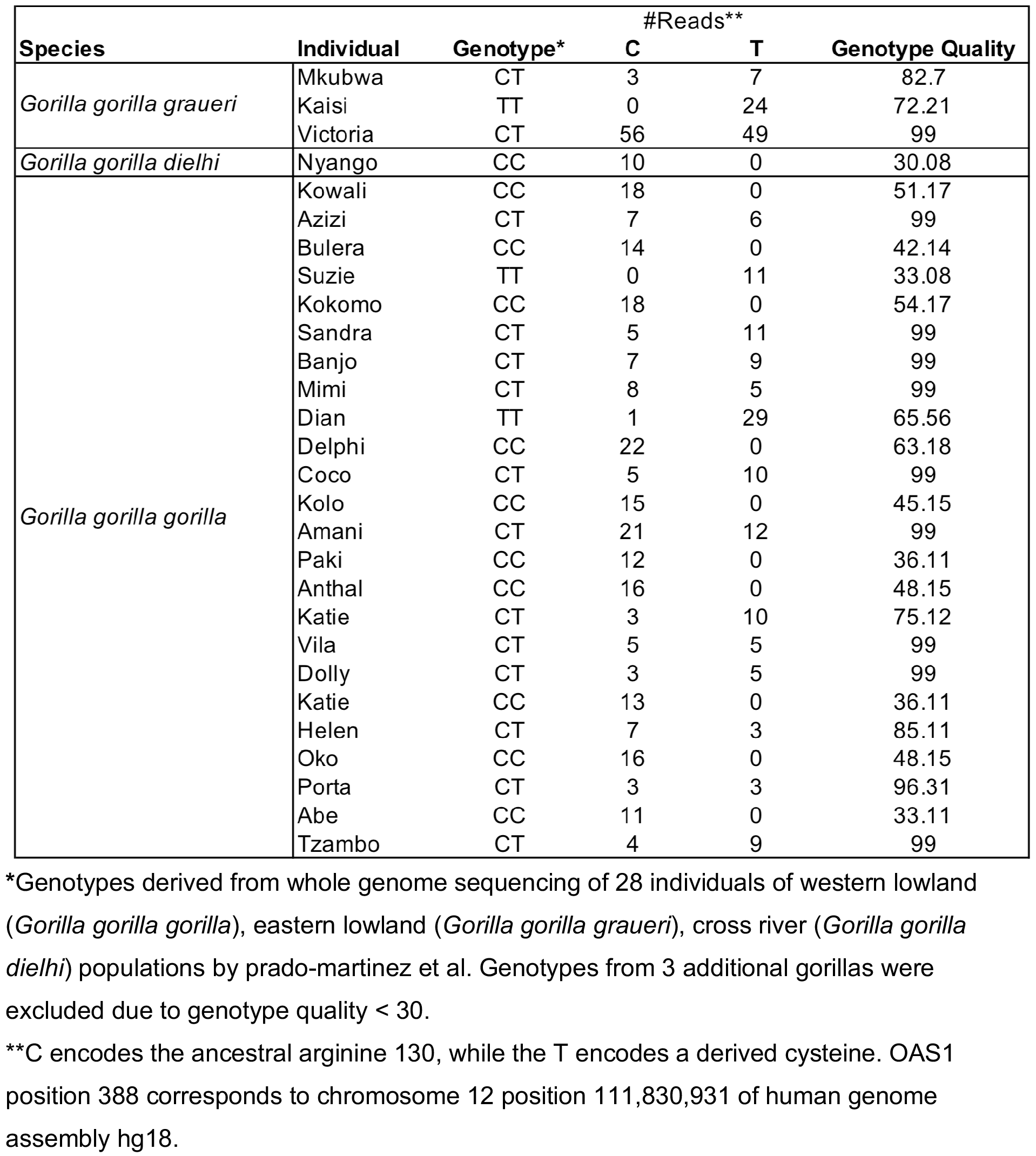
Distribution of gorilla genotypes at OAS1 nucleotide 388.

